# The Impact of Low T3 Syndrome on Lipid Metabolism and Gut-Brain axis in Patients with Bisphenol A-Associated Intracerebral Hemorrhage

**DOI:** 10.64898/2026.05.07.723656

**Authors:** Guoping Wang, Jinhua Chen, Zhongdong Qiao, Dongxing Guo, Ping Guo, Aiven Wang, Wanli Sun, Ji-yuan Lyu

**Author notes:** Corresponding author;Guoping Wang.

## Abstract

**BACKGROUNG:** Bisphenol A (BPA) has been linked to hypertension and disturbances in lipid metabolism; however, limited evidence is available regarding its association with hypertensive intracerebral hemorrhage (ICH).

**METHODS:** A multicenter, retrospective case-control study was conducted involving 129 participants, including individuals from an ICH group and healthy controls. Standard assays were employed to assess serum thyroid function, lipid profiles, serum fatty acid-binding ※protein 4 (FABP4), oxidative stress markers, gap junction proteins, Wnt/β-catenin signaling pathway activity, and expression changes of S100A8-mediated inflammatory cytokines involved in gut-brain interactions. Correlation analyses using Pearson and Spearman methods revealed that both BPA exposure and low T3 levels were significantly associated with elevated diastolic blood pressure, altered lipid metabolism, gut microbiota composition, and microglial activation.

**RESULTS:** Gender-based disparities in lipid metabolism were identified. Changes in β3-adrenergic receptor and neuromodulin-1 expression appear to influence fat regulation and attenuate oxidative stress responses. Subsequently, increased expression of gap junction proteins and activation of the Wnt/β-catenin signaling pathway contribute to metabolic reprogramming and alterations in biochemical kinetics. Gut microbiota analysis demonstrated that, compared to controls, the ICH group exhibited significant dysbiosis and reduced alpha diversity. Further correlation analyses indicated that BPA levels were positively associated with FABP4 and oxidative stress markers, while S100A8 showed a strong dependence on microglial expression.

**CONCLUSION:** The interplay between lipid metabolism dysfunction and pro-inflammatory cytokines enhances vascular vulnerability. Collectively, BPA exposure, oxidative stress, and microglia-mediated neuroinflammation are significantly associated with an elevated risk of hypertensive ICH.

**China Clinical Trial Registry registration notice:** From: China Clinical Trials Registry +To:guopingwang60a yunyanshuangfei

**FUNDING:** This work was supported by the Natural Science Foundation of Shanxi Province (grant no. 201701D121177)

**Key information:** Gender-specific differences were observed in lipid metabolism and oxidative stress parameters; BPA exposure was shown to induce lipid metabolic disturbances, promote excessive production of oxidative stress byproducts, and consequently elevate oxidative stress responses; BPA was associated with stress-induced alterations in thyroid hormone function, further exacerbating dysregulation of lipid metabolism and oxidative stress; Fatty acid binding protein 4 (FABP4), a key adipokine implicated in metabolic disorders and adipose tissue inflammation, exhibited a significant positive correlation with serum BPA levels, whereas low levels of triiodothyronine (T3) were negatively correlated with FABP4. These findings suggest that serum FABP4 may serve as a biochemical marker for chronic low-grade adipose tissue inflammation and metabolic dysfunction; Gap junction proteins and the Wnt/β-catenin signaling pathway may contribute to microglial activation and mediate neuroinflammatory responses, nerve injury, and secondary pathological processes in obesity-related cerebral hemorrhage.

## Introduction

Intracerebral hemorrhage (ICH) accounts for approximately 10% to 15% of all stroke cases. The primary etiological factors include common genetic variations and vascular risk factors, such as hypertension and blood pressure variability(1-3). Recently, the evaluation of risk factors associated with ICH pathogenesis has become a focal point of research (4). One emerging question is whether long-term exposure to bisphenol compounds may serve as a potential risk factor for ICH. Given that bisphenols are recognized as widespread environmental toxins, investigating their peripheral genotoxic effects, immune cell dynamics, and association with stroke risk may help elucidate the detrimental impacts of lipid metabolism disorders and oxidative stress responses on neuronal cells and large vascular functions. The Wnt/β-catenin signaling pathway plays a regulatory role in metabolic functions and exerts physiological protective effects. Dysregulation of this pathway has been linked to metabolic disorders and cerebrovascular diseases (5). Abnormal or multidirectional blood flow can promote endothelial dysfunction and is closely associated with atherosclerosis (6). Therefore, a potential correlation exists between lipotoxicity and the protective Wnt/β-catenin signaling pathway. Changes in the pathophysiological state, such as the downregulation of connexin 43 (Cx43), may provide insights into the disruption of normal cerebrovascular homeostasis and function (7).

Bisphenol A (BPA) contains endocrine-disrupting properties that interfere with thyroid function and contribute to obesity, resulting in an imbalance in the intestinal microbiota and significant alterations in intestinal biomarkers related to alpha diversity. The microbial community’s richness demonstrates a positive correlation with Clostridium and Salmonella, while showing a negative correlation with Enterobacteriaceae, Shigella, Bifidobacterium, and Lactobacillus (8). Animal studies have confirmed that BPA is associated with disruptions in tissue metabolic pathways (9). Furthermore, research indicates that BPA can significantly elevate the level of oxidative stress (OS) response in tissue cells. It is well established that β3-adrenergic signaling plays a role in regulating adipocyte anabolism, and low levels of triiodothyronine amplify cyclic adenosine monophosphate (cAMP) signaling, potentially leading to disturbances in lipid metabolism (10). Lipotoxicity and neurocytotoxicity interact with multiple organ functional mechanisms, possibly involving protein receptor pathways, immune system activity, and epigenetic mechanisms underlying inflammatory response (11). Superoxide anion (O_2_^−^), a reactive oxygen species (ROS), is generated during pathophysiological processes and acts as a mediator oxidant, reflecting the overall intracellular ROS levels (12). The elevation of tissue O_2_^−^ levels interacts with the activity of immune-inflammatory cytokines and contributes to neurotoxicity (13). The paraventricular nucleus of the hypothalamus modulates sympathetic nerve output in response to stress and chronic regulatory stimuli, thereby inducing vasosensitive chronic hypertension mediated by α2δ-1-associated N-methyl-D-aspartate receptors within the paraventricular nucleus (14). This process ultimately facilitates the development of intracerebral hemorrhage (ICH) following cerebrovascular injury. Intestinal microbiota may influence both the immune system and cerebrovascular function through the production of metabolites and neurotransmitters. Alterations in the population of intestinal neurons exhibiting immune responsiveness to neuregulin 1 (NRG1) may attenuate the oxidative stress (OS) response, indicating a neuroprotective effect (15). In addition, astrocytes release functional mitochondria into neighboring cells to support the maintenance of brain cellular homeostasis and to exert protective effects against oxidative stress and inflammatory responses (16).Mitochondrial dysfunction and its regulatory mechanisms represent a key focus in research on the pathogenesis of intracerebral hemorrhage (ICH). The critical involvement of mitochondrial dysfunction in the progression of arterial hypertension (AH), atherosclerosis, and post-hemorrhagic brain injury is now widely acknowledged (17).

## Methods

### Study population

The clinical data of patients diagnosed with intracerebral hemorrhage (ICH) and admitted to multiple hospitals between February 2016 and April 2019 were collected. 129 individuals with a long-term history of exposure to plastic products were identified. The age range of these subjects was 18 to 51 years, with a mean age of 38 ± 8.3 years.Among the selected cases, 65 patients were categorized into the low free triiodothyronine (LT3) group based on FT3 levels (The LT3 standard and gender subgroups are presented in the supplementary file -7 “Method 1”.).

Data collection was conducted through experimental research and clinical observation, both of which adhered to the standards established by the American Stroke Association, the Cardiovascular and Stroke Care Committee, and the Clinical Neurology Committee (18). Obesity was diagnosed according to the World Health Organization (WHO) criteria for the Asia-Pacific region: normal weight (BMI < 25 kg/m^2^) and obese (BMI ≥ 25 kg/m^2^)(19). Hypertension was defined as systolic blood pressure ≥140 mmHg and/or diastolic blood pressure ≥90 mmHg (1 mmHg = 0.133 kPa)(20). Diabetes diagnosis followed the 2022 (21)(24hABP, Follow-up visits, Selection criteria, and The positive control groups,Exclusion criteria are presented in the supplementary file -7 ‘methods’ 2,3,4,5).

### Routine detection

Serum levels of testosterone and thyroid hormones were measured using electrochemiluminescence immunoassay, along with inflammatory markers, neuropeptide Y, and bisphenol A. The serum oxidative enzyme system was assessed via the enzyme rate method, and the release of superoxide anion (O_2_^−^) in serum was determined using the cytochrome C reduction assay. Brain tissue samples were processed to culture approximately 2 g of brain cells. Following cell attachment, the DCFH-DA probe was applied for in situ detection of reactive oxygen species by fluorescence microscopy. Intracellular superoxide anion levels were evaluated using DHE staining, while mitochondrial superoxide anion levels were measured using MitoSOX Red fluorescence. Serum concentrations of fatty acid binding protein 4, S100A9, S100A8, glial fibrillary acidic protein, and TMEM119 were quantified using a double-antibody sandwich enzyme-linked immunosorbent assay (ELISA). (Routine detection are presented in the supplementary file -7 ‘methods’ 6)

### Western blot and Cell culture

Intestinal Neuregulin1 (NRG1), brain tissue NRG1, Connexin 43 (CX43), Yes associated protein (YAP), Wnt and β-catenin; Western blotting and real-time quantitative PCR (RT-PCR) were used to detect changes in protein levels. Cell culture: The specific steps are as follows; (Western blot and Cell culture are presented in the supplementary file -7 ‘methods 7,8).

**Serum metabolomics analysis**, **Intestinal Content Sequencing Sequencing and Bioinformatics Analysis of Intestinal Flora (**are presented in the supplementary file -7 ‘methods’ 9,10,11).

### Statistical analysis

General demographic data are presented as continuous variables with mean ± SD. Normality tests are conducted for indicators, and the T-test is employed to compare the means of two small samples. For qualitative data comparison, one-way ANOVA and Chi-square tests are utilized. The differences in serum BPA concentration, biomarkers, and intestinal flora are analyzed by Spearman correlation data, processed with R software, and a heatmap is applied. The independent sample Mann-Withney-Wilcoxon test is used for the analysis of bacterial alpha diversity among different groups and for the analysis of differences between the data of the case group and the control group. Non-parametric tests are also employed for the analysis of significant differences in the relative abundance of various bacterial groups at the generic level. The combination and cross-effect of serum BPA, CX43, YAP, and intestinal flora expression levels are analyzed by the Logistic regression model, and the relative risk ratio, attribution ratio, and interaction index are calculated. Statistical analysis is performed using SPSS v20.0 software (SPSS Inc, USA), and a difference is considered statistically significant (P<0.05).

## Results

**FIG 1**.. Characteristics of lipid metabolism and oxidative stress products in patients with low T3 and SCH and gender differences between groups. (**a**) Compared with the control group, the level of BPA in LT3 and SCH groups was significantly increased, and MG was significantly higher than FMG, with gender differences. (**b**) Compared with the control group, serum APO-β, apo CIII, APO-β /apoA1 ratio, BMI, HOMA-IR, TG, FBG, LDL-C and Tch in LT3 and SCH groups were significantly increased, while HDL-C was significantly decreased; Compared with FMG group, APO-β, apo CIII and LDL-C of MG patients in LT3 and SCH groups were increased, but LT3-MG group was significantly increased. Compared with MG group, BMI and TG of FMG patients in LT3 and SCH group showed an increasing trend. apo-β was significantly increased in LT3-b group compared with LT3-a group. It indicates that there are significant gender differences in blood lipid metabolism. (**c**) FT3 in LT3 group was significantly lower than that in control group; However, FT3 decreased more significantly in LT3-b group than in LT3-a group. TSH in SCH group increased significantly compared with control group. The proportion of women is obviously greater than that of men. These results indicate that ICH has different stress thyroid function status, showing different sex ratio phenomenon. (**d**) Compared with the control group, the serum levels of 3-nitrotyrosine and malondialdehyde (MDA) in ICH patients were significantly increased, while the level of superoxide dismutase (SOD) was significantly decreased; In addition, compared with LT3-FMG, the increase of 3-nitrotyrosine was more obvious in LT3-MG patients, indicating that oxidative stress products also had gender differences. (**e**) Compared with the control group, nDBP, DBP and NPY were significantly increased in both LT3 and SCH groups. (**f**) Compared with the control group, serum inflammatory cytokines IL-1β, TRL-4, NF-kBP56 and MYD88 were significantly increased in LT3 and SCH groups. (**g**) Compared with the control group, serum superoxide anion (O^2-^) and Hydrogen peroxide (H2O2) concentration in LT3 and SCH groups was significantly increased. (**h**)Compared with the control group, serum levels of fatty acid binding protein 4 (FABP4),S100A8,and S100A9—as well as the anti-transmembrane protein 119 antibody (anti-TMEM119 antibody), a well-established marker of microglial stability—were significantly elevated in both the LT3 and SCH groups (all P < 0.05).

**Figure 1.**
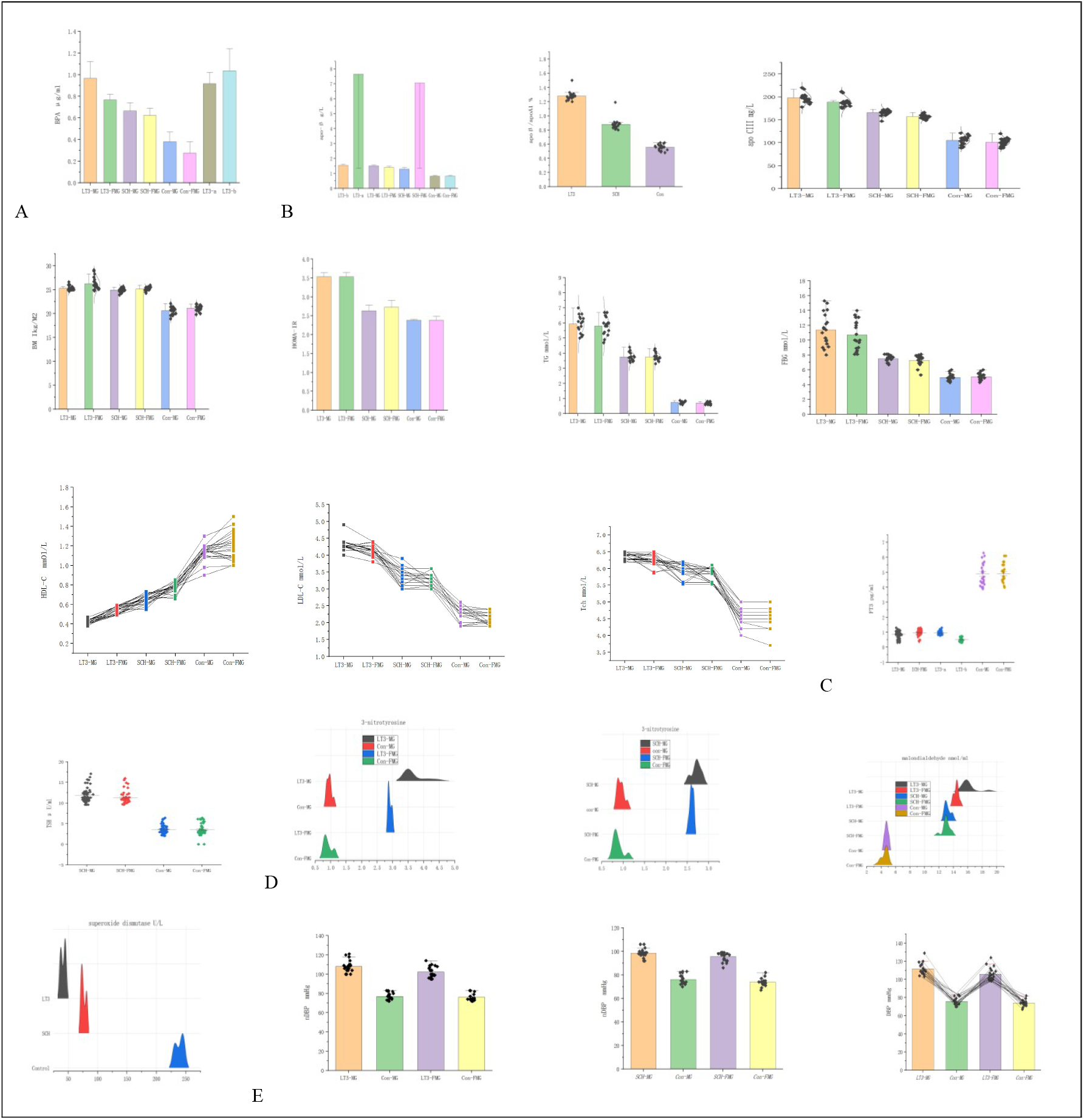
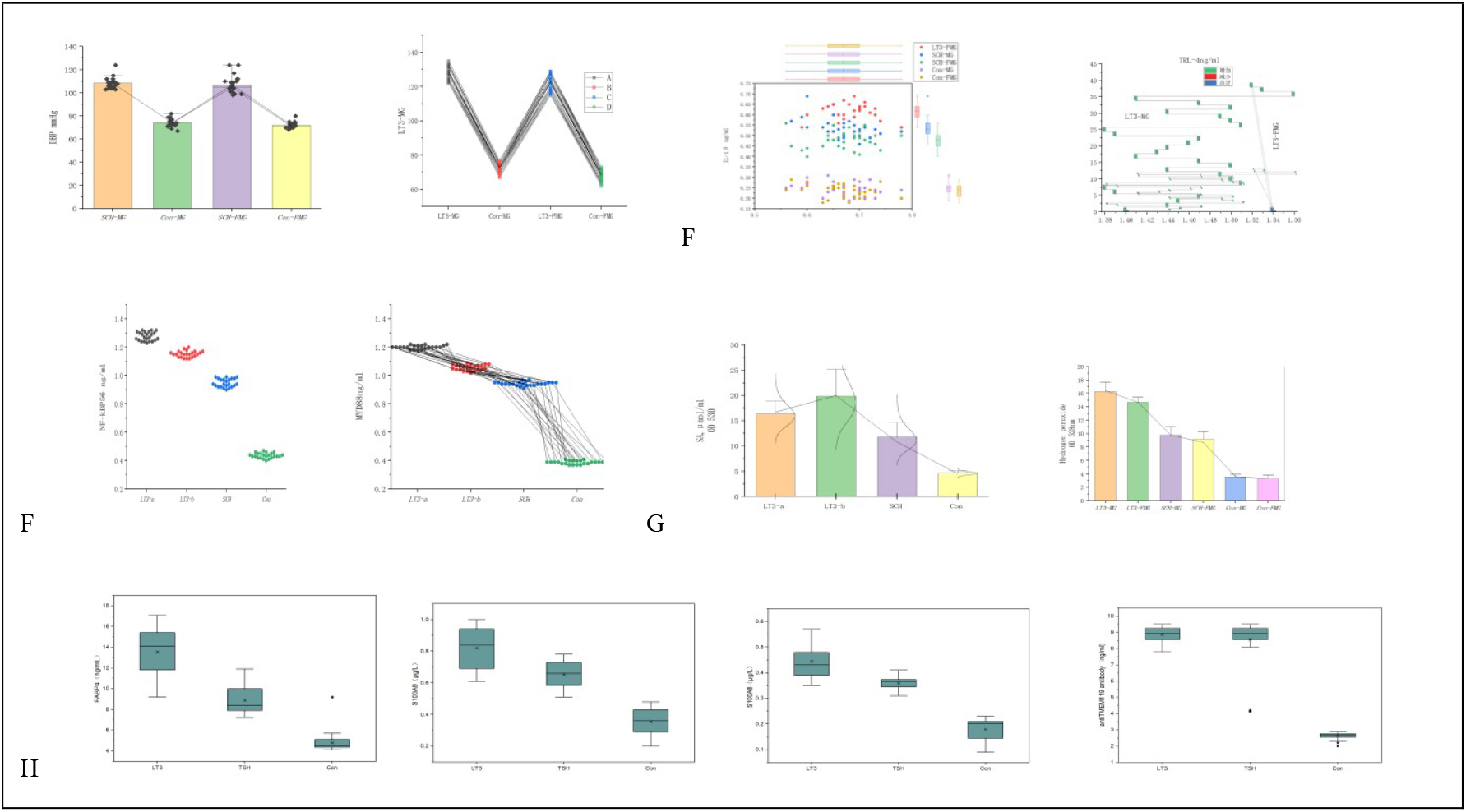
Characteristics of serum lipid metabolism and oxidative stress products in patients with low T3 and gender differences between groups. *P < 0.05, ** P < 0.01, between LT3 and control; †P < 0.05, ††P < 0.01, between SCH and control ; ‡ P < 0.05, ‡ ‡P < 0.01. between MG and FMG. BPA, bisphenol-A,; apo-β,lipoprotein β; apo CIII, lipoprotein CIII; apoβ/apoA1ratio; BMI, body mass index; HOMA-IR, Homeostatic model assessment of insulin resistance; TG, Triglycerides.; FPG, fasting plasma glucose; HDL-C, high density lipoprotein cholesterol; LDL-C, low density lipoprotein cholesterol;Tch, Total cholesterol;nDBP, nightime diastolic.blood pressure; DBP, nightime diastolicblood pressure;Neuropeptide Y,NPY; 3-nitrotyrosine;MDA,malondialdehyde;SOD,superoxide dismutase; IL--1β, Interleukin-1β;TRL-4,toll-like receptor-4,NF-kBP56,nuclear factor-kBP56,MYD88, myeloiddifferentiation factor 88; SA, (O^2-^) Superoxide anion; H2O2,Hydrogen peroxide.

**FIG. 2**. Comparison of total superoxide and lipid oxidative stress messenger protein expression between groups. (**a**)Compared with SCH group and positive control group, brain tissue reactive oxygen species (ROS) increased significantly in LT3 group. However, LT3-MG-b and LT3-FMG-b subgroups were significantly increased compared with LT3-MG-a and LT3-FMG-a, with statistical significance. (**b**) The expression of β3-AR and UCP1 in LT3 group was decreased compared with the control group and SCH group, and the decrease was more significant in the male group compared with the female group. (**c**) Compared with LT3 group, NRG-1 expression in intestinal and brain tissues of SCH group was more significantly increased; Compared with MG group, NRG-1 increased in FMG group. (**d**) Compared with LT3 group, CX43 expression increased more significantly in SCH group. However, the expression of CX43 showed temporal differences, and the expression level decreased with the increase of time. (**e**): Compared with SCH group, the expression of LT3 group was significantly reduced.. (**f**) Compared with the control group and the LT3 group, both men and women showed a significant increase in β-connexin in the SCH group. (**g**) Wnt 1 levels in LT3-MG, LT3-FMG and SCH-MG were significantly lower than those in the control group. (**h**) Compared with the positive control group, the intracellular and mitochondrial superoxide anion concentration and mitochondrial hydrogen peroxide level in LT3 and SCH groups were significantly increased. All the above parameters were P < 0.05.

**Figure 2.**
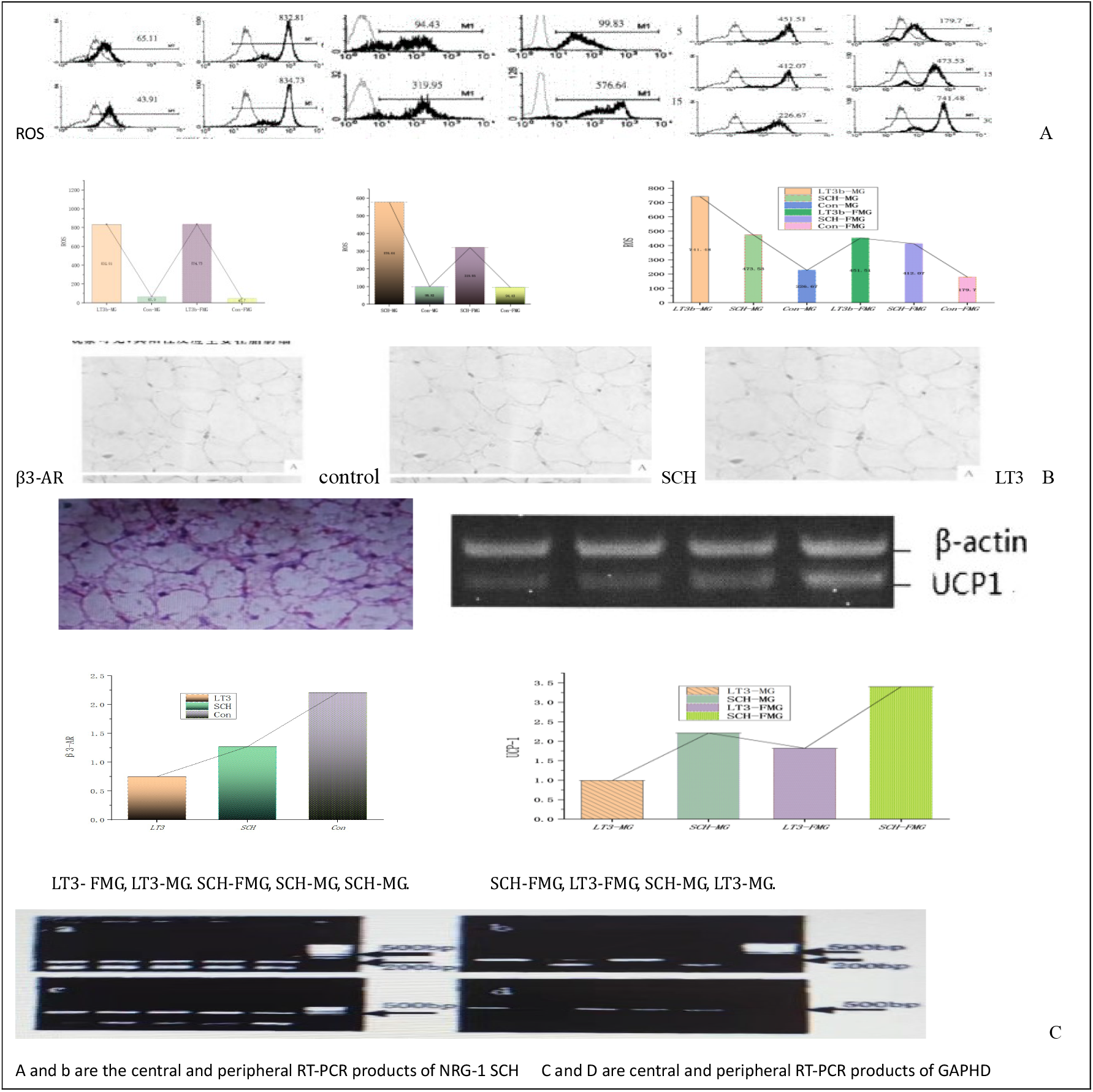
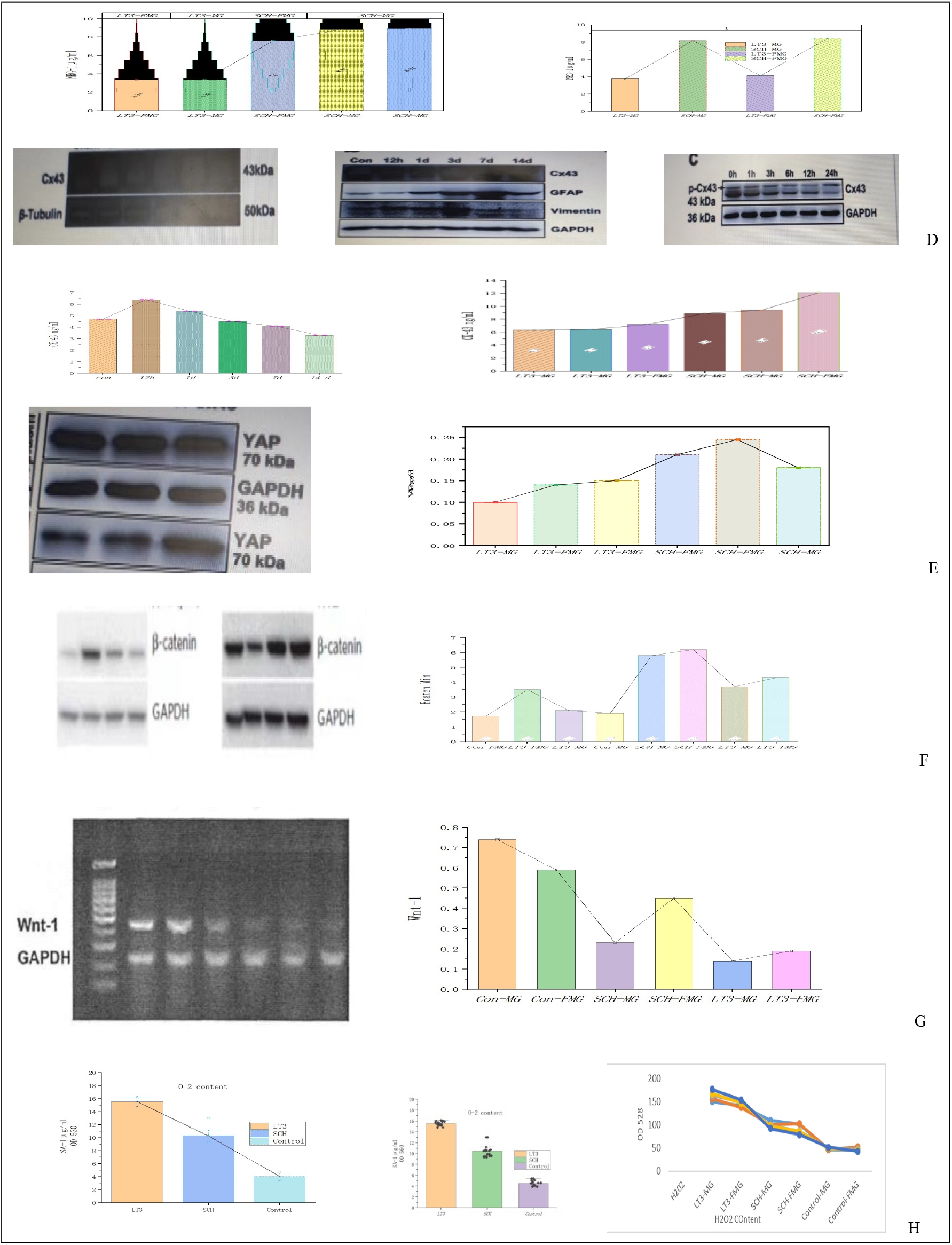
Comparison of total superoxide and lipid oxidative stress messenger protein expression among all groups. *P < 0.05, ** P < 0.01, between LT3 and control; †P < 0.05, ††P < 0.01, between SCH and control ; ‡ P < 0.05, ‡ ‡P < 0.01. between MG and FMG. ROS, Reactive oxygen species; MG, Male group; FMG, Female group; β3-AR, β3-adrenergic receptor; UCP-1,Uncoupling proteins; NRG1,Neuregulin1;CX43, Connexin 43 ; YAP, Yes associated protein ; SA, (O^2^-), superoxide anion.

**FIG 3** (**a**) At the phylum level, Firmicutes, Bacteroidetes, Actinobacteria and Proteobacteria accounted for more than 99% of the flora. Compared with the control group, the number of Firmicutes in LT3-a, LT3-b and SCH groups decreased ; Compared with the control group, Bacteroidetes increased in LT3-a, LT3-b and SCH groups ; Compared with the control group, Actinomycetes decreased in LT3-a, LT3-b and SCH groups (8.18%vs5.8%vs12.3%vs19.2%)P=2.64×10^−-6^, Mann-Whitney U test; Compared with the control group, Proteobacteria increased in LT3-a, LT3-b and SCH groups. (**b**)Genus level. Compared with the control group, Haldemanella, Ruminococcus torques group and paraprevotella had the largest microbial populations in LT3 and SCH groups. (**c**) Compared with the control group, the least number of microorganisms in LT3 and SCH groups were Parabacteroides, Roseburia, Bifidobacteria, LachnospiraceaeUCG008, Actinomyces and Lactobacillus.To detect differences between the three groups, Linear discriminant analysis Effect Size (LEFSe) was used for a detailed quantitative analysis of the relative abundance of bacterial communities at the genus level among the different groups.At the generic level, according to the difference in the relative abundance of characteristic bacteria (relative abundance > 1%, P < 0.05). (**d**) Compared with the control group, the blood pressure-related bacteria in LT3 and SCH groups were Eubacterium limosum and Streptococcus, and the insulin resistence-related markers Prevotella-9 and Paraprevotella.Escherichia Castellani and Shigella were significantly increased in oxidative stress-related bacteria. (**e**) Compared with LT3 and SCH, the control group showed significant increases in Chromobacterium, Rothella, Erysipelothrix, and Prevotella 2 indicating differences in strains between SCH and LT3 groups and the control group. (**f**) At the genus level, the relative abundances of Bacteroides fragilis, Romboutsia and Ruminant -5 were statistically significant between SCH and LT3 groups. (Discriminant Analysis Canonical discriminant analysis Classify, summarize the training sample data and the Beta diversity Index are presented in the supplementary file -8 Reesults 1)

**Figure 3.**
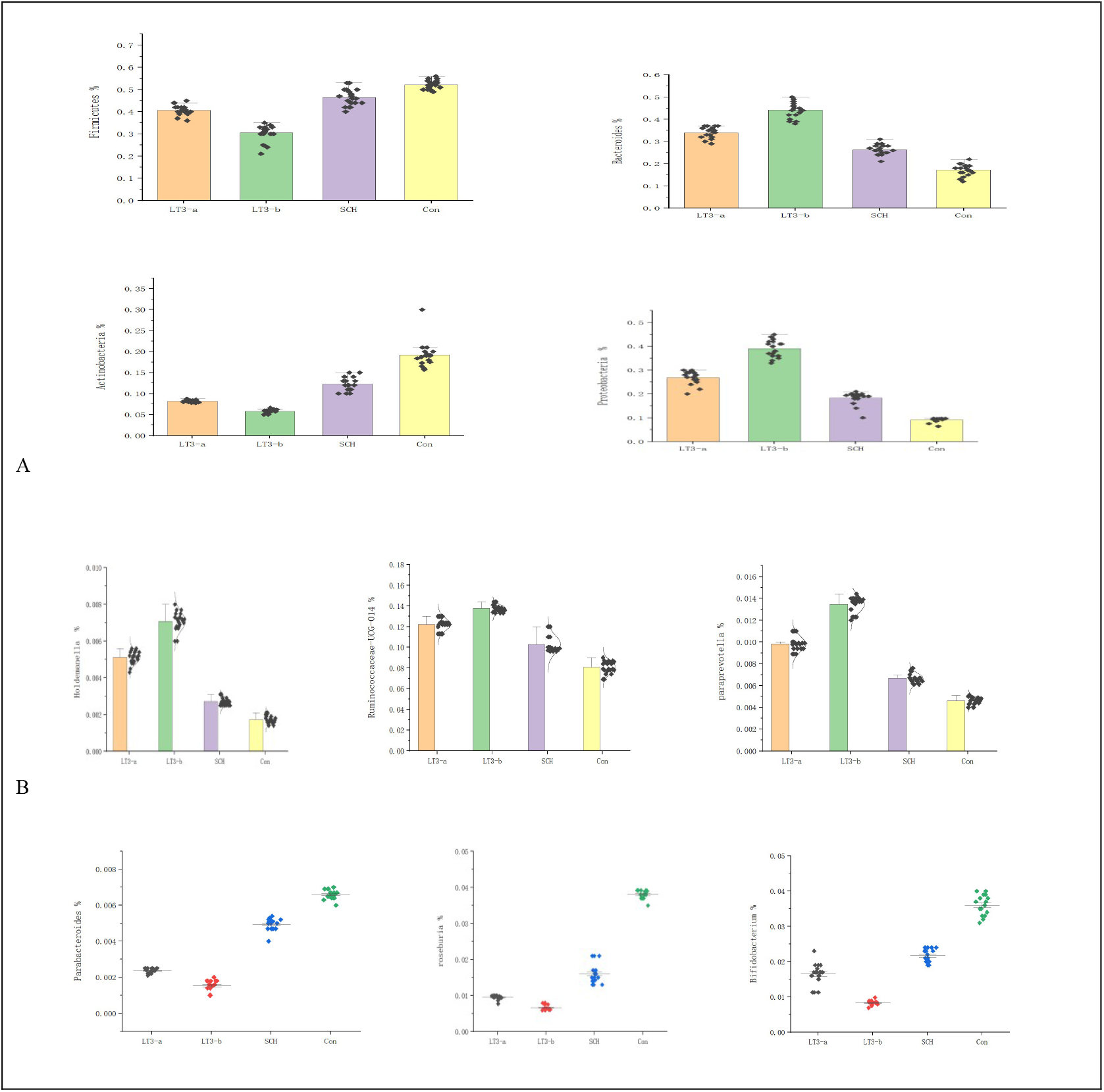
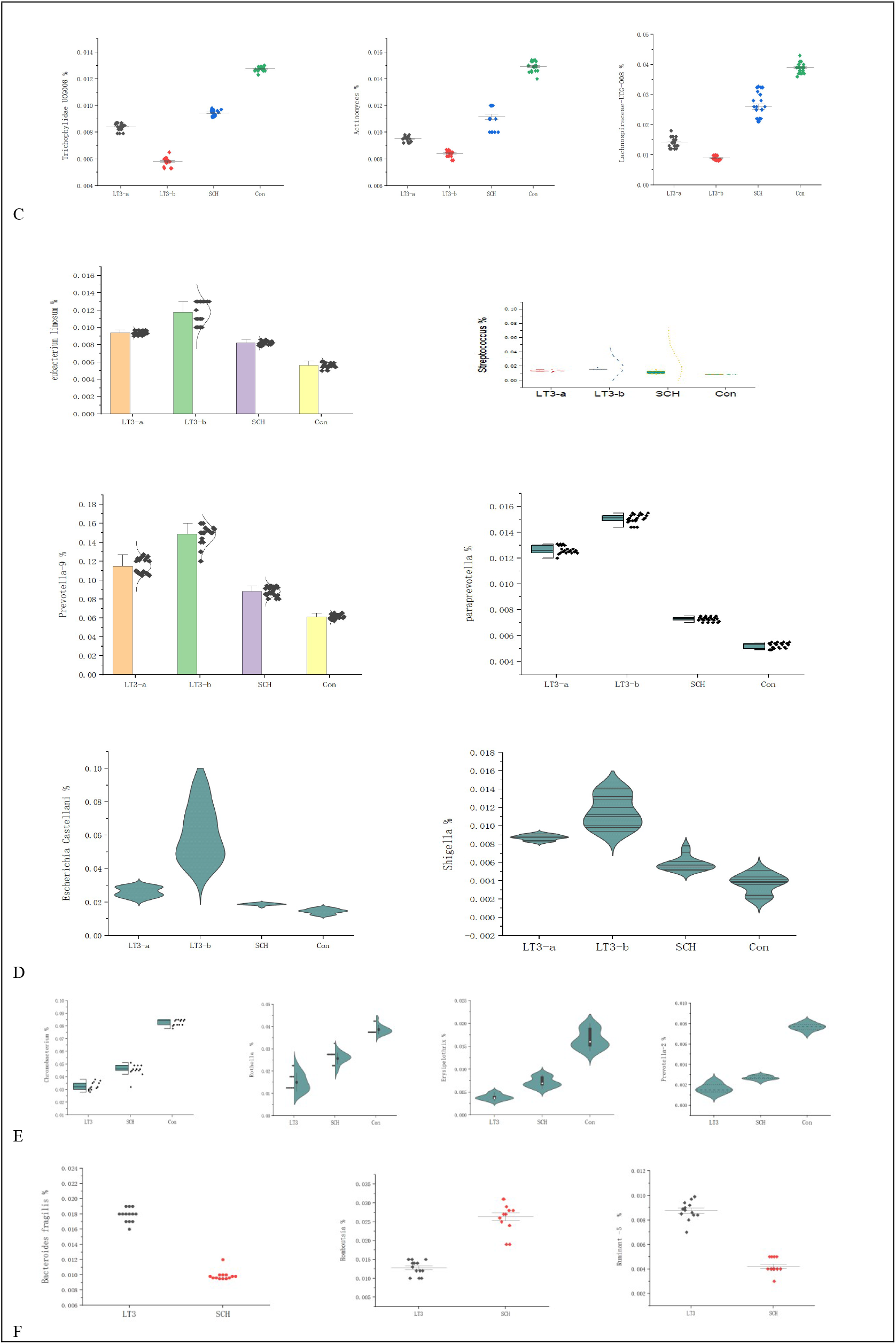
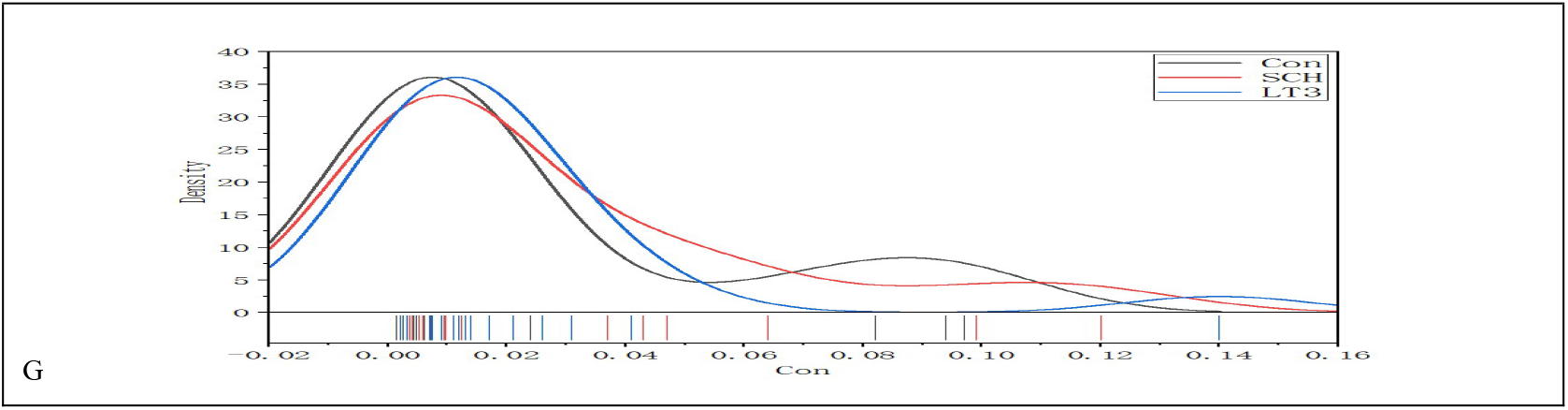
Comparison of intestinal flora between groups. *P < 0.05, ** P < 0.01, between LT3 and control; †P < 0.05, ††P < 0.01, between SCH and control ; ‡ P < 0.05, ‡ ‡P < 0.01. between MG and FMG. FMG, the female group; MG, the male group; SCH, Subclinical hypothyroxinemia T3, Low T3 syndrome.

**FIG 4 Correlation analysis between clinical indicators and intestinal flora (** Correlation analysis and intestinal flora are presented in the supplementary file -8 Reesults 2

**Figure 4.**
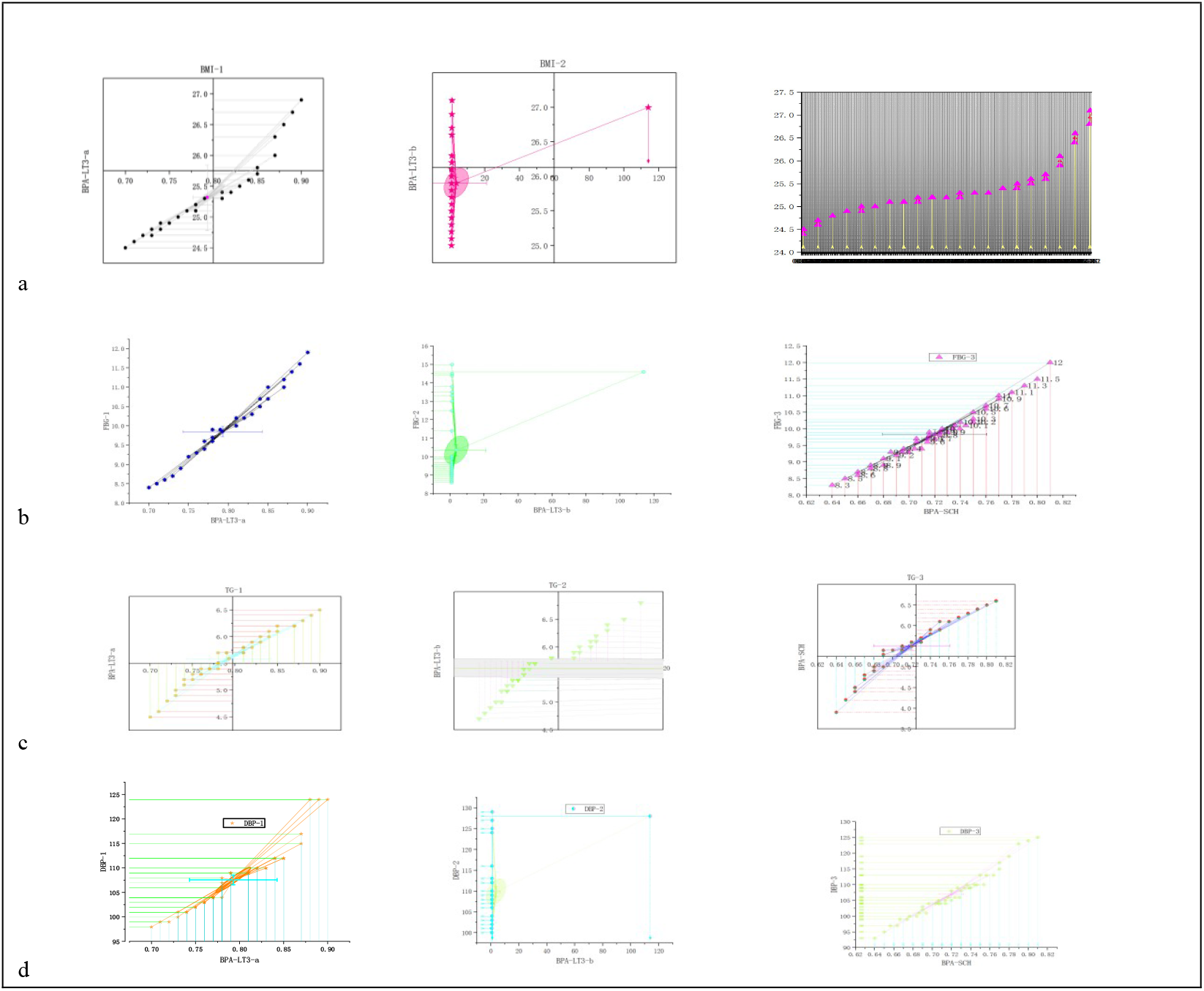
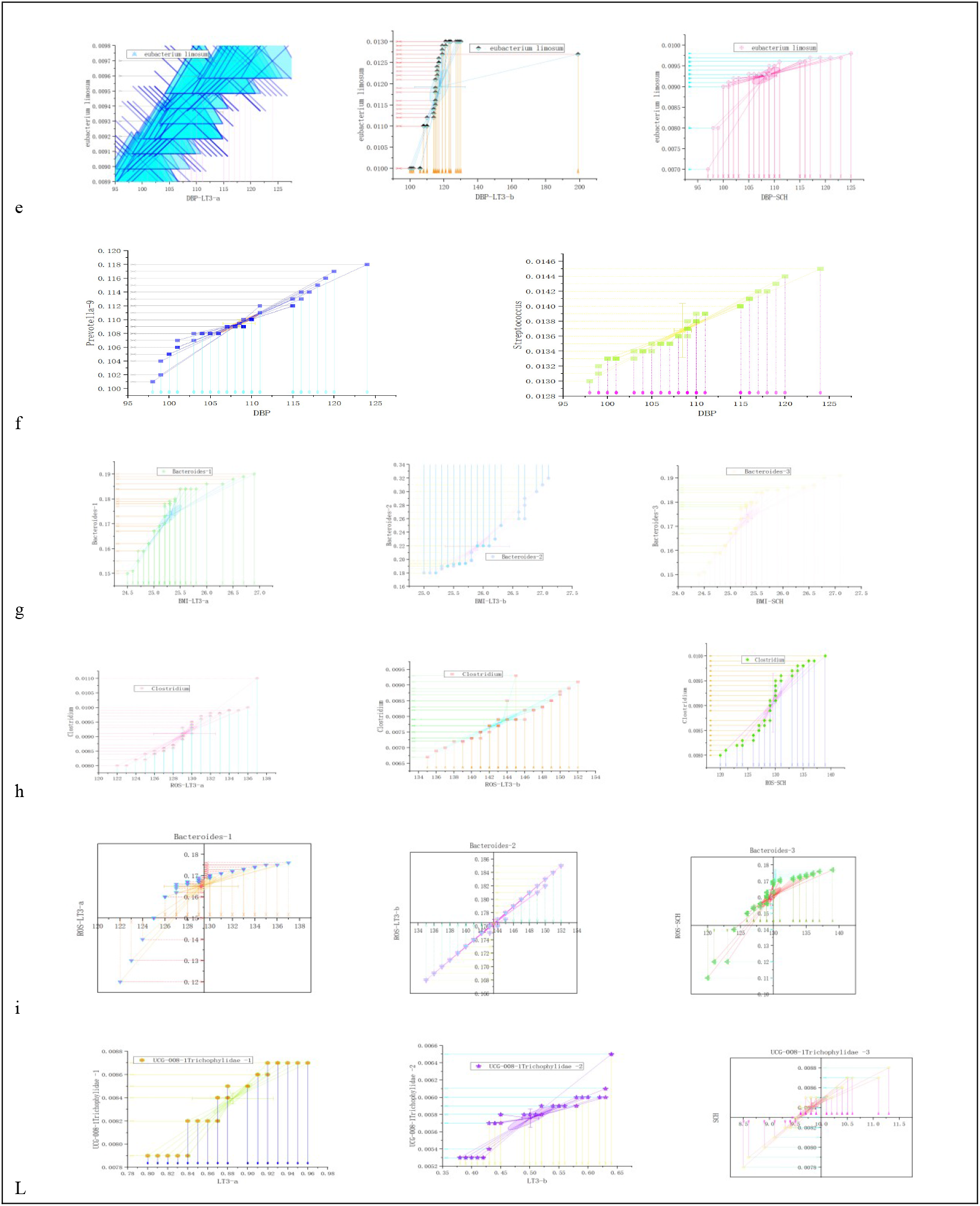
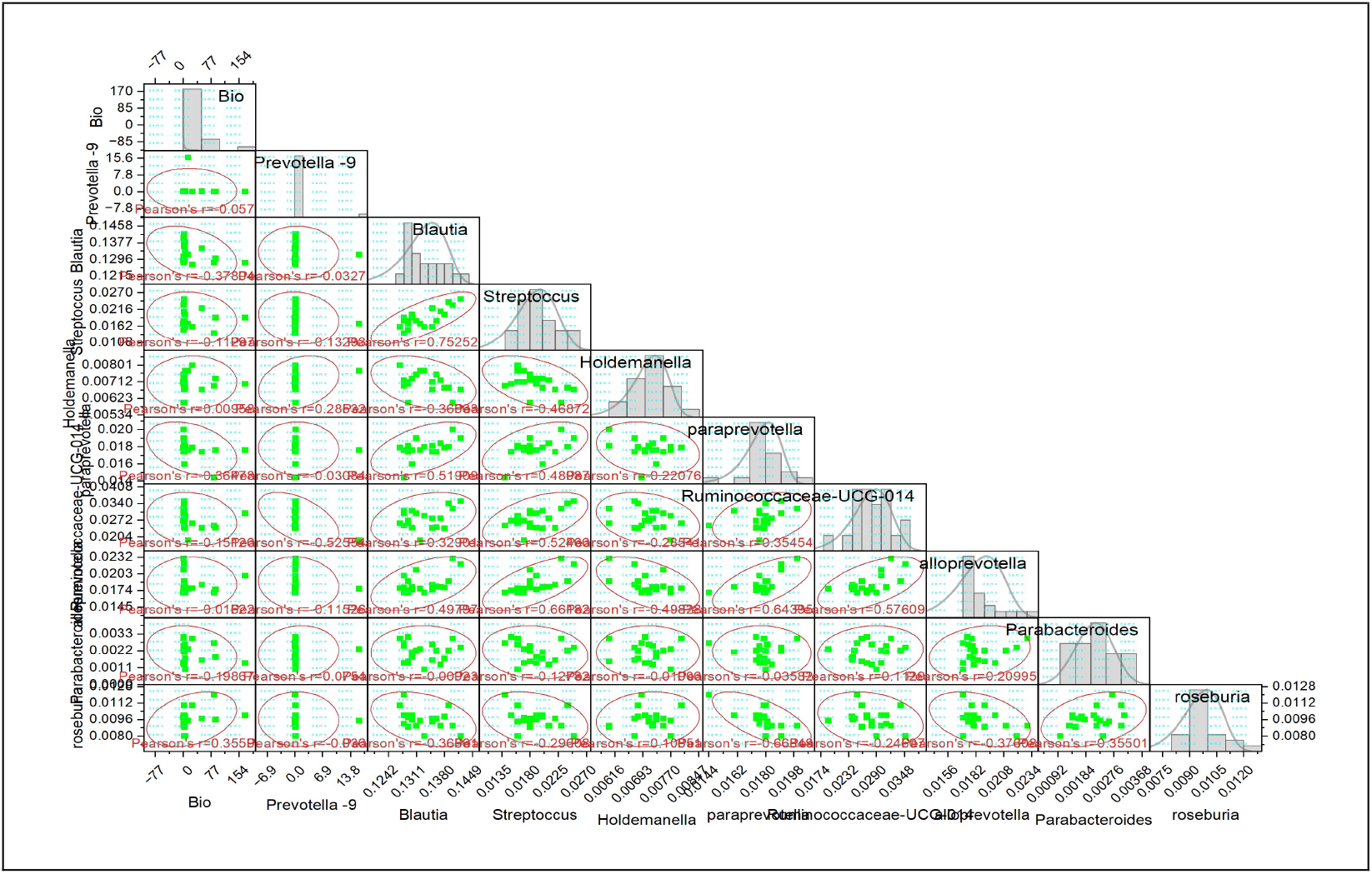
Correlation analysis between clinical indicators and intestinal flora. *P < 0.05, ** P < 0.01, between LT3 and control; †P < 0.05, ††P < 0.01, between SCH and control; ‡ P < 0.05, ‡ ‡P < 0.01. between MG and FMG..*P < 0.05, ** P < 0.01, between bio and intestinal flora; SCH, Subclinical hypothyroxinemia T3, Low T3 syndrome.Bio,Biomarker;BPA, Bisphenol-A; T3, Low T3 syndrome; TG,Triglyceride; LDL-C,Low-density lipoprotein cholesterol; HDL-C, High-density lipoprotein cholesterol;Tch,Total cholesterol; apoβ,Apolipoproteinβ; ApoA1, Apolipoprotein A1;apo C III,Apolipoprotein CIII;apoE,Apolipoprotein E;ROS,reactive oxygen species;MDA, Malondialdehyde; SOD,Superoxide; DismutaseCh/pl,ACholesterol/Phospholipids;DA,Adenosine deaminase; DPP-4, Dipeptidyl peptidase-4.

**FIG 5 Heat map and correlation analysis of serum metabolomics and microbiota abundance clustering analysis** (a,b,c are presented in the supplementary file -8 Reesults 3)

**Figure 5.**
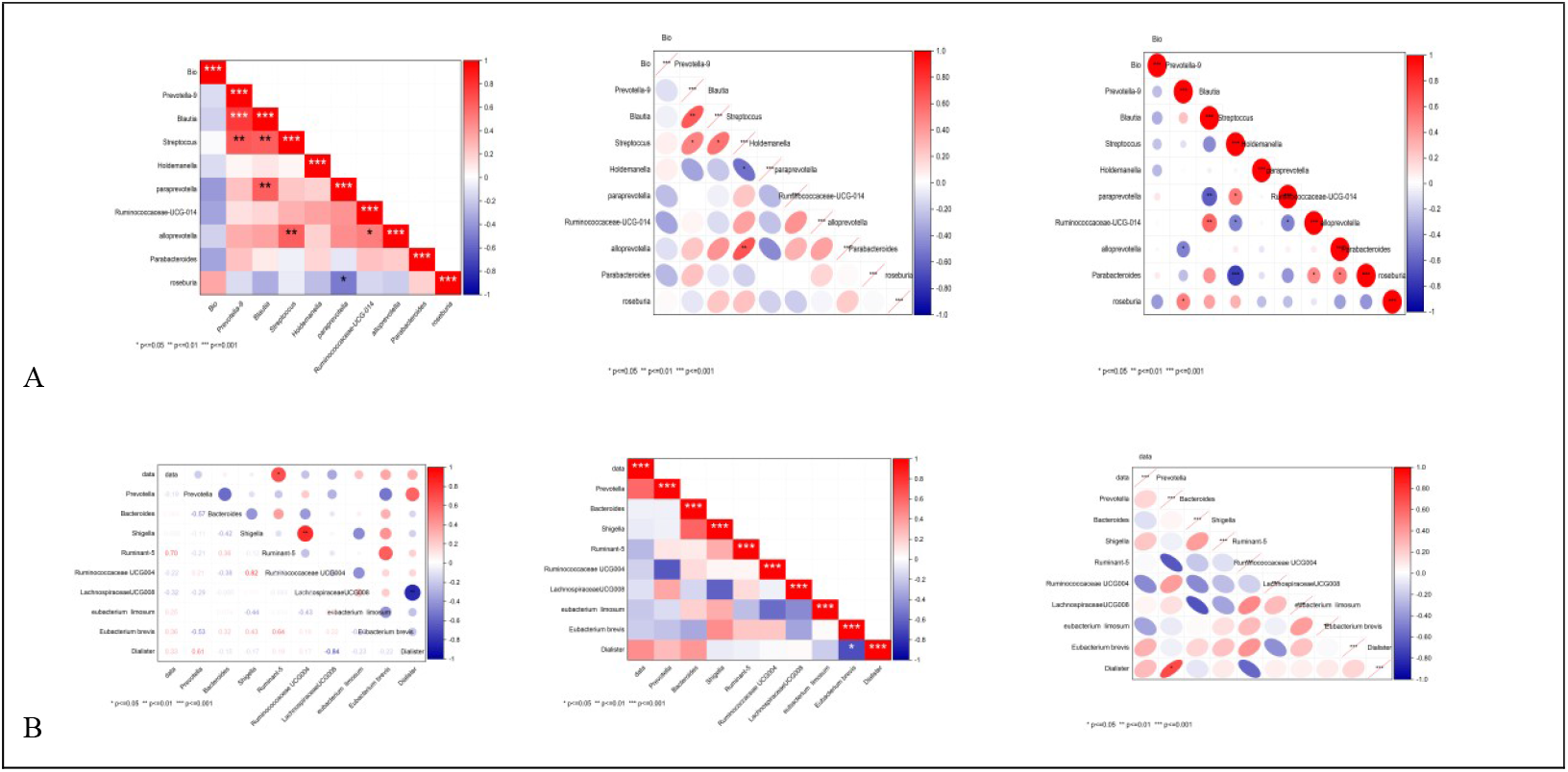
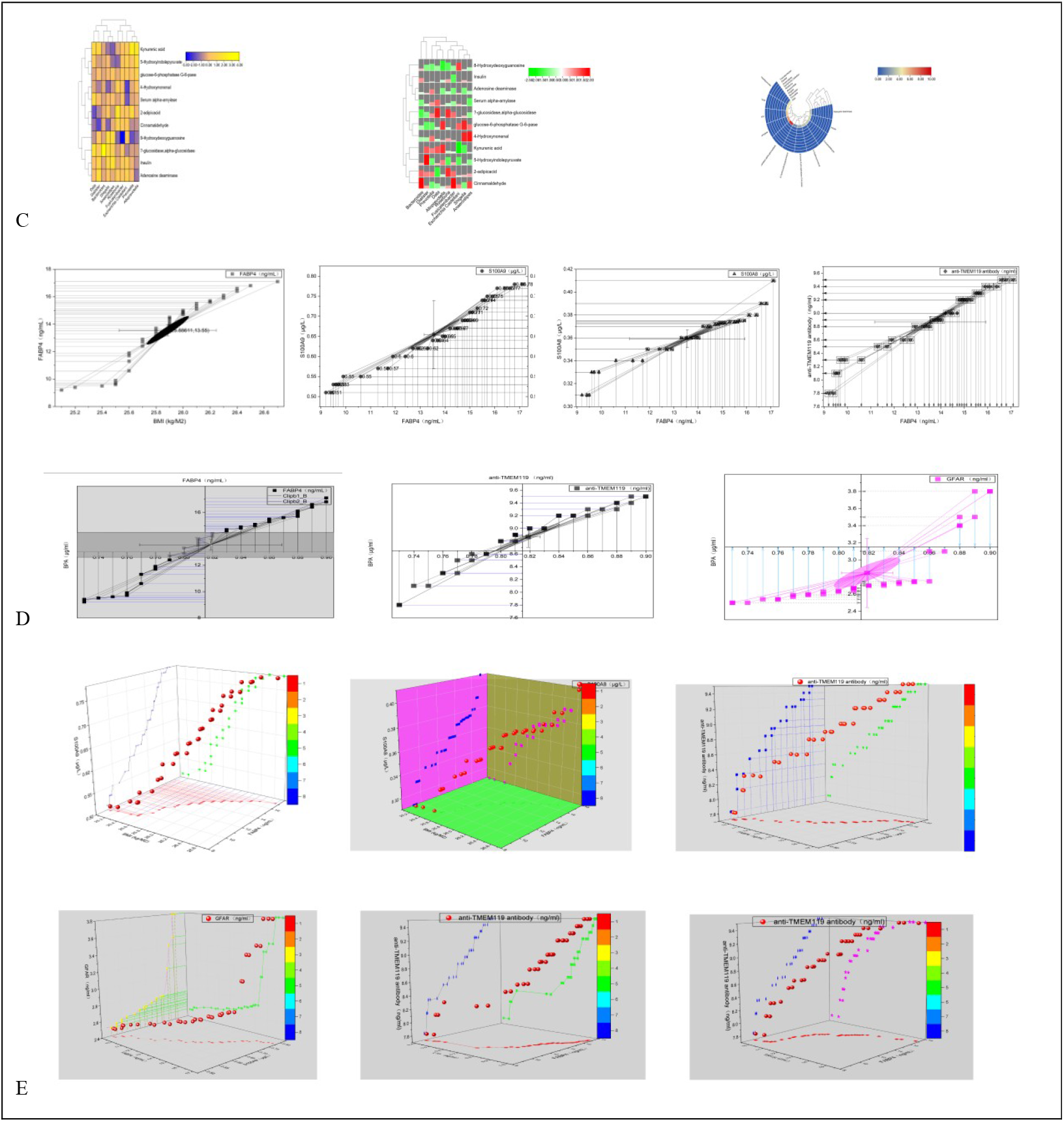
Heat map and correlation analysis of serum metabolomics and microbiota abundance clustering analysis. *P < 0.05, ** P < 0.01, between LT3 and control; †P < 0.05, ††P < 0.01, between SCH and control ; ‡ P < 0.05, ‡ ‡P < 0.01. between MG and FMG. FMG, the female group; MG, the male group; SCH, Subclinical hypothyroxinemia T3, Low T3 syndrome; Apo-E. lipoprotein-E; UDP-glucose, Uridine 5’-diphosphoglucose; TG, triglyceride; ROS, Reactive oxygen species; LDL-C, low density lipoprotein-Cholesterol; Apo-A1, lipoprotein A1; MDA, malonaldehyde; SOD, Superoxide Dismutase; apoβ, lipoprotein β; HDL-C, High density lipoprotein-Cholesterol; Apo-CIII, lipoprotein-CIII; Ch/pl, cholesterol/ phospholipids; FABP4,fatty acid binding protein 4; BPA,bisphenol A; GFAP,glial fibrillary acidic protein; TMEM119,Transmembrane Cell 119

**FIG 6** (**a**) Representative CT images reveal hyperdense lesions localized to the thalamus and basal ganglia in both the LT3 and SCH groups of patients with cerebral hemorrhage. (**b**) Quantitative RT-PCR analysis demonstrates a statistically significant upregulation of β-catenin expression in the LT3 group relative to the SCH group. (**c**) Immunohistochemical assessment indicates a marked increase in microglial density in both the LT3 and SCH groups compared with the positive control group. (**d**) While the proportion of third-generation astrocytes and oligodendrocytes exhibits a downward trend in the LT3 and SCH groups versus the positive control group, immunofluorescence co-localization analysis reveals significantly enhanced expression specifically at the direct interface between oligodendrocyte progenitor cells and astrocytes (all P < 0.05)..

**Figure 6.**
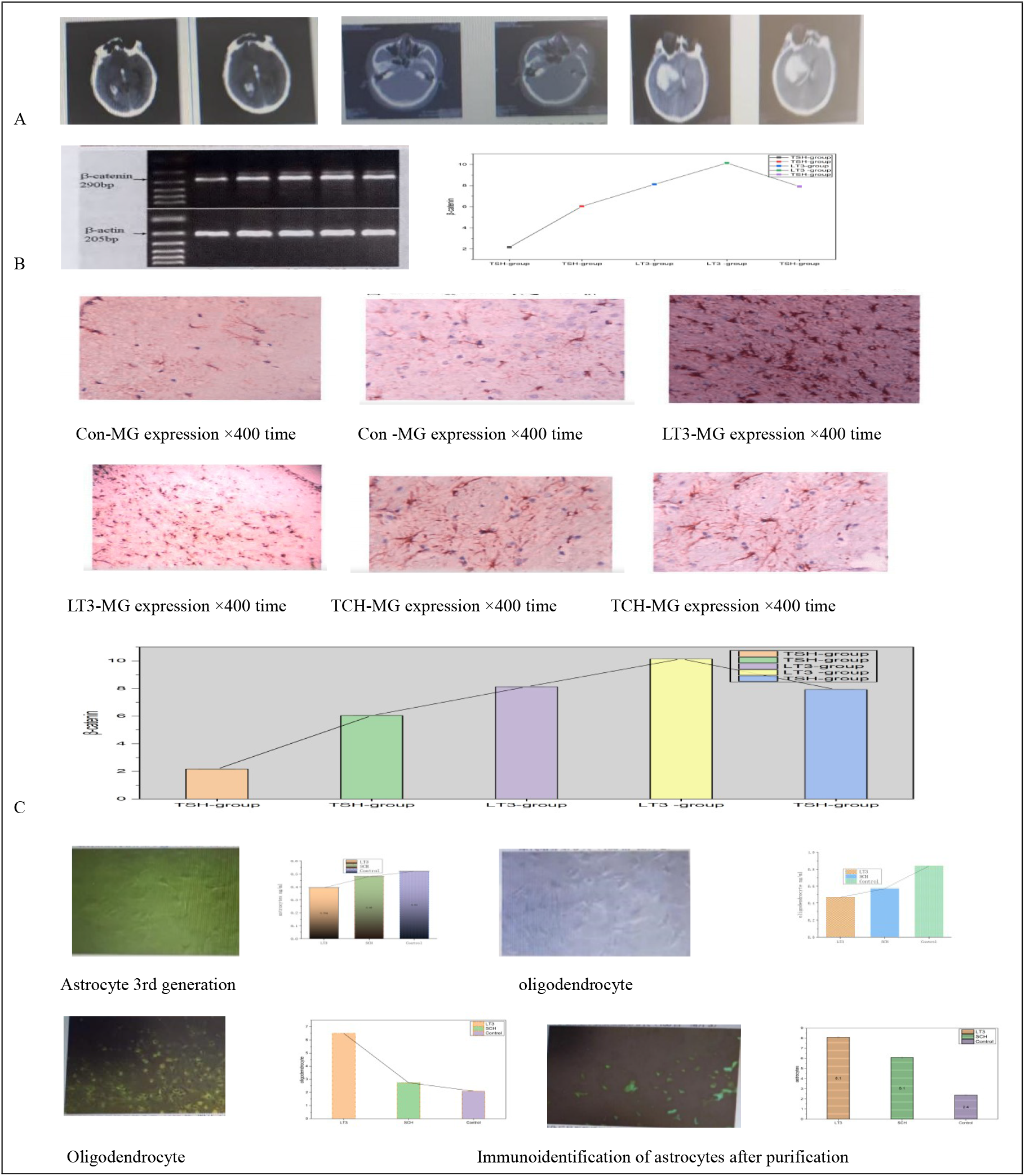
CT images of ICH patients, showing the absence of the winged (Wnt)/β-catenin signal transduction pathway in brain tissue, and the expression of microglia and astrocytes. *P < 0.05, ** P < 0.01, between LT3 and control; †P < 0.05, ††P < 0.01, between SCH and control; ‡ P < 0.05, ‡ ‡P < 0.01. between MG and FMG. MG, microglia;Con,Control; SCH, Subclinical hypothyroxinemia T3, Low T3 syndrome

### Relationship between BPA and risk factors of OS expression

(BPA risk factors expression and logistic regression of risk factors are presented in the supplementary file -8 Reesults 5, 6).

## Discussion

Bisphenol A (BPA) is a well-documented environmental endocrine-disrupting chemical implicated in the pathogenesis of obesity and metabolic disorders through mechanisms aligned with the “environmental obesogen hypothesis. (10)” Epidemiological evidence consistently demonstrates a significant positive association between urinary or serum BPA concentrations and the incidence of hypertension (22). Recent mechanistic studies have elucidated that BPA exposure disrupts key metabolic pathways—particularly those governing glucose homeostasis, lipid metabolism, and redox balance—via dual actions: (i) estrogen receptor–mediated endocrine disruption and (ii) transcriptional and epigenetic reprogramming of metabolic genes. These findings establish a systems toxicology framework for evaluating the metabolic health risks posed by persistent environmental pollutants (11, 23). Specifically, BPA downregulates mRNA expression of critical regulators of glucose metabolism—including glucokinase (GCK), insulin-like growth factor 1 (IGF-1), and glucose transporter 2 (GLUT2)—while concurrently upregulating adaptor protein 1 (AP1), thereby activating downstream lipid metabolism genes. This coordinated dysregulation positions BPA-sensitive transcripts as potential biomarkers of impaired insulin signaling and glucose utilization (12, 24, 25). In murine models, BPA exposure induces oxidative stress gene expression, impairs mitochondrial fatty acid oxidation and amino acid catabolism, and triggers gut microbiota–associated ecological shifts—culminating in adipose tissue expansion and systemic metabolic inflammation (26, 27). Furthermore, BPA–driven oxidative stress and lipid dyshomeostasis exacerbate obesity-related comorbidities, including metabolic inflammation and atherosclerosis progression, implicating redox-sensitive pathways—such as HDL oxidation, β-catenin signaling interference, and superoxide anion–mediated endothelial dysfunction—as novel therapeutic targets (28-30). Notably, excessive accumulation of BPA and structurally analogous compounds promotes vascular endothelial dysfunction and dysregulates Wnt/β-catenin signaling in brain tissue, contributing to neuroendocrine impairment (31, 32).

Emerging data further indicate that BPA modulates intestinal microbial ecology: human and animal studies report reduced alpha diversity and altered taxonomic composition, including decreased abundance of *Bifidobacterium*, *Lactobacillus*, and *Actinomyces*, alongside enrichment of *Ruminococcaceae* UCG-004 and *Ruminococcus* group 5 (Refs. 10, 12, and present study). Importantly, these shifts correlate with host metabolic phenotypes—e.g., *Ruminococcus*-UCG-014 depletion is positively associated with colonic short-chain fatty acid levels (Ref. 12). Given the intestine’s role as a central hub for thyroid hormone (TH) metabolism—regulated by TH transporters (e.g., MCT8, OATP1C1), activating/deactivating enzymes (e.g., DIO2/DIO3), and nuclear receptors (e.g., TRα1)—BPA–induced gut dysbiosis may perturb the intestinal-thyroid axis, thereby influencing systemic TH bioavailability and action.

At the molecular level, connexin 43 (Cx43), a gap junction protein with antioxidant regulatory functions, interacts with the transcriptional co-activator Yes-associated protein (YAP). Cx43 downregulation promotes YAP nuclear translocation, modulating glutamate and ion homeostasis, cholesterol/sphingolipid metabolism, and inflammatory responses to environmental stressors (33). Concurrently, TLR4/NF-κB signaling is amplified—a pathway known to mediate post-hemorrhagic neuroinflammation, immune cell polarization, and oxidative damage following intracerebral hemorrhage (ICH)(34–36). Thus, targeting Cx43/YAP dynamics and TLR4–NF-κB crosstalk represents a promising strategy for mitigating ICH-associated secondary injury (37).

Our clinical and preclinical analyses reveal robust gender- and thyroid-status–stratified patterns in BPA-associated cerebrovascular pathology. Patients with low triiodothyronine (LT3) syndrome and subclinical hypothyroidism (SCH) exhibited significantly elevated serum BPA, apolipoprotein B, apolipoprotein CIII, and LDL-C—particularly in males—alongside heightened 3-nitrotyrosine (a marker of nitrative stress) in male LT3 patients. Conversely, females demonstrated higher BMI and triglyceride levels, and a stronger inverse correlation between serum lipoprotein (a) [Lp (a)] and low T3 levels—suggesting sex-specific modulation of lipid metabolism under hypothyroid conditions. Moreover, among SCH patients, females showed disproportionately elevated TSH levels compared to males. Brain tissue analyses revealed markedly increased ROS levels in the LT3 group versus controls and SCH; correspondingly, β3-adrenergic receptor (ADRB3) and uncoupling protein 1 (UCP1) expression were suppressed—most prominently in males. Intestinal and cerebral neuregulin-1 (NRG1) expression was significantly upregulated in the SCH group relative to LT3, especially in females, whereas Cx43 expression was most profoundly diminished in the LT3 group. Mitochondrial and cytosolic superoxide anion and hydrogen peroxide levels were also significantly elevated across LT3 cohorts.

Microbiome profiling confirmed distinct dysbiosis signatures: both LT3 and SCH groups exhibited depletion of beneficial taxa (*Bifidobacterium*, *Lactobacillus*, *Actinomyces*) and expansion of *Ruminococcaceae* UCG-004 and Ruminococcus group 5, without concomitant increases in Enterobacteriaceae or *Lactobacillus* (contrary to prior reports). PCoA at the phylum level revealed >99% representation by Firmicutes, Bacteroidetes, Proteobacteria, and Actinobacteria; LT3 and SCH groups shared parallel trends—decreased Firmicutes and Actinobacteria, increased Bacteroidetes and Proteobacteria—relative to controls. At the genus level, dominant taxa included *Holdemania*, *Ruminococcus gnavus*, and *Paraprevotella*, while *Parabacteroides*, *Roseburia*, *Bifidobacterium*, *Erysipelotrichaceae* UCG-008, *Actinobacteria*, and *Lactobacillus* were consistently depleted. Correlation analyses identified clinically relevant associations: diastolic blood pressure correlated positively with Firmicutes abundance and *Streptococcus*; BMI correlated with baseline *Bacteroides* levels; and plasma BPA concentration correlated positively with HOMA-IR and *Holdemania* abundance.

Collectively, these findings demonstrate that BPA–induced dysregulation of lipid metabolism, low-grade inflammation, and gut microbiota imbalance converge to activate the Wnt/β-catenin signaling axis, thereby coordinating S100A8 expression, microglial activation, and astrocytic reactivity. This tripartite interaction establishes a neurovascular-inflammatory nexus central to the pathophysiology of BPA–associated cerebral hemorrhage—providing mechanistic insight into its inflammatory etiology and identifying actionable molecular and microbial targets for intervention.

## Acknowledgments

This work was supported by the Natural Science Foundation of Shanxi Province (Grant No. 201701D121177). The authors would like to extend their sincere gratitude to Dr. Zong Liang from the Third Clinical College of Changzhi Medical University for his technical support. Additionally, the research team gratefully acknowledges the group of senior experts and doctoral students from Shanghai Jiao Tong University for their valuable suggestions and critical comments on the manuscript.

---

**Table 1.**
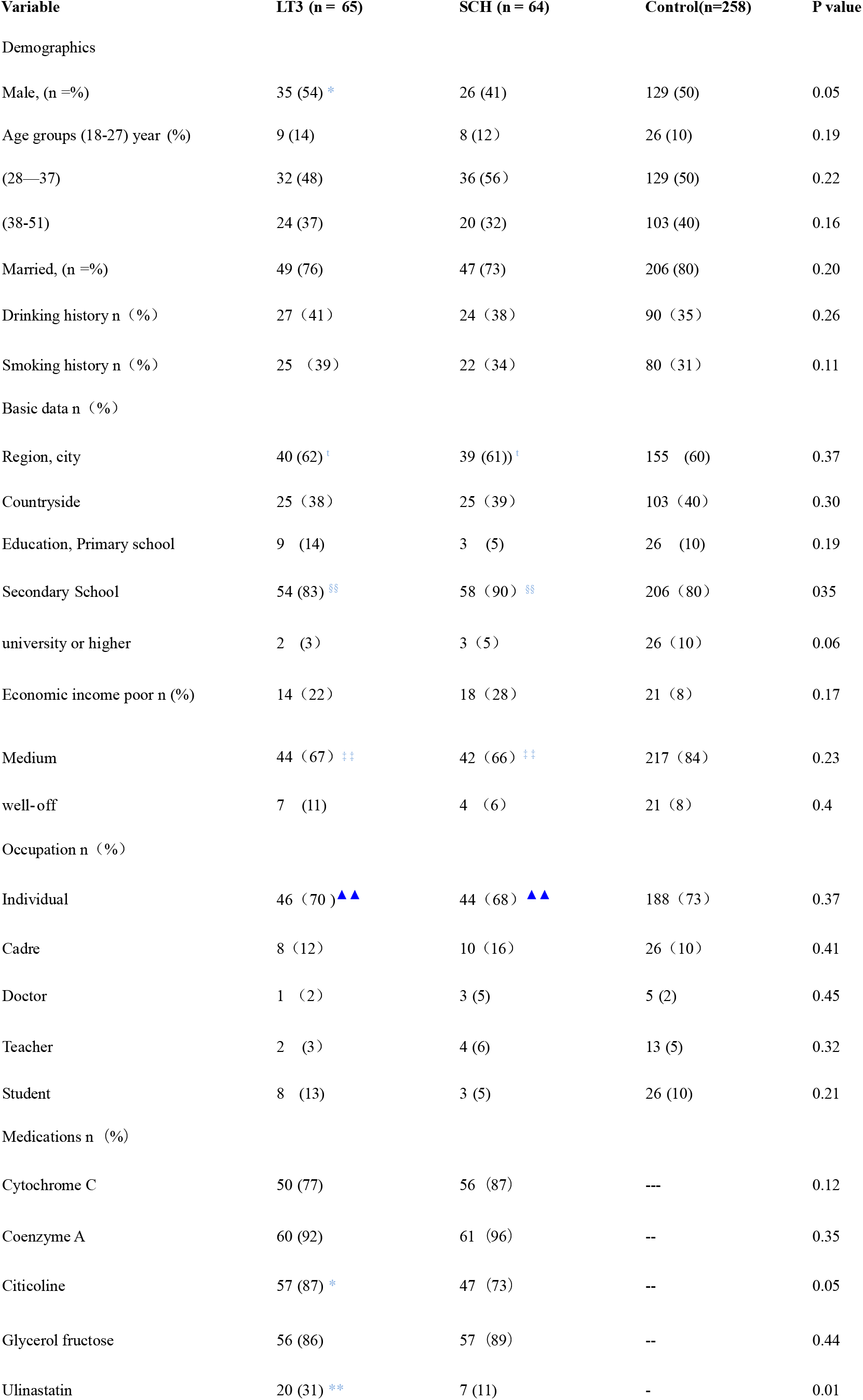
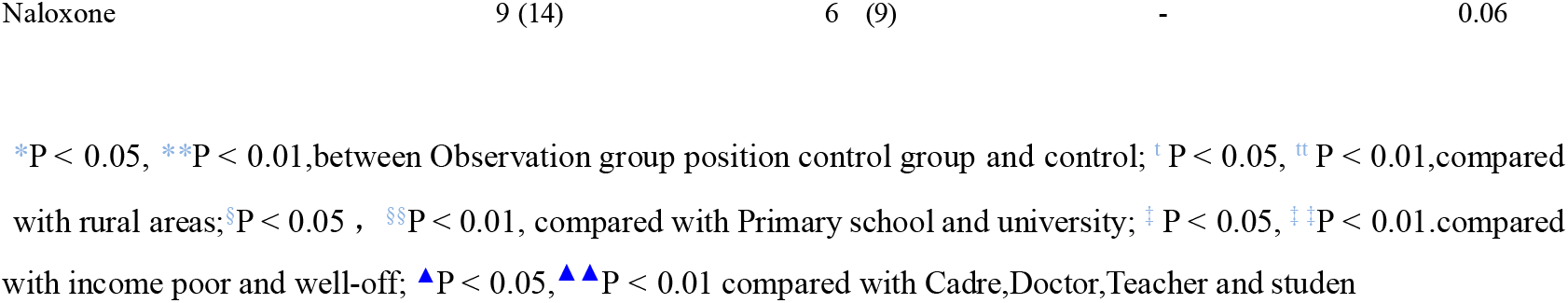
Characteristics of the study population.

**Table 2.** Alpha diversity index among different groups.

| Parameter | LT3 | SCH | Control | P value |
| --- | --- | --- | --- | --- |
| Observed-species | 493.6±21.5** | 544.566±21.4† | 842.1±20.6 | 0.02 |
| shannon | 4.9±1.05** | 5.56±1.5† | 10.15±1.7 | 0.01 |
| simpson | 0.799±0.06**‡ | 0.931±0.12†† | 1.69±0.2 | 0.01 |
| chaol | 451.03±34.4* | 501.9±33.2 | 776.11±13.2 | 0.05 |
| ACE | 542.43±26.15* | 593.2.82±31.05 | 923.24±28.1 | 0.05 |
| Goods-coverage | 104.55±16.6 | 118.73±15.2 | 184.4±12.1 | 0.52 |
| PD-whole-tree | 19.84±5.51** | 24.03±5.37 † | 42.14±6.19 | 0.04 |

\*P < 0.05, \*\* P < 0.01, between LT3 and control; †P < 0.05, ††P < 0.01, between SCH and control; ‡ P < 0.05, ‡ ‡P < 0.01. between LT3 and SCH.

